# Therapeutic efficacy of hiMSC-derived Extracellular Vesicles from Serum-containing and Xeno-free media for osteoarthritis treatment

**DOI:** 10.1101/2025.08.20.671255

**Authors:** S. Sana Sayedipour, Jelle Nikkels, Tobias Tertel, Helena E. D. Suchiman, Matilde Balbi, Georgina Shaw, Luis J. Cruz, Louise van der Weerd, Chiara Gentili, Bernd Giebel, Josephine Mary Murphy, Ingrid Meulenbelt, Yolande F. M. Ramos

## Abstract

**Background:** Extracellular vesicles derived from human induced mesenchymal stromal cells (hiEVs) constitute a promising cell-free therapeutic option for osteoarthritis. To allow translation to the clinic we evaluated the therapeutic effects of hiEVs for osteoarthritis treatment. Specifically, we assessed the efficacy of hiEVs collected from serum-containing and serum-free, PurStem (PS), media in an OA mice model.

**Methods:** hiEVs were administered with or without hydrogel via intra-articular (i.a.) injection in a destabilization of the medial meniscus (DMM) mouse model. Fluorescence imaging was used to monitor the retention of IR780-labeled hiEVs in the joint cavity. The therapeutic effects were evaluated by analyzing damage scores as well as catabolic and anabolic markers, including Mmp13 and Col2 expression, in joint tissues.

**Results:** Fluorescence imaging confirmed that hiEVs remained localized at the injection site without systemic migration. HiEVs demonstrated significant protective effects against joint tissue degeneration in the DMM mouse OA model, as evidenced by reduced damage scores, decreased Mmp13 expression, and increased anabolic processes (Col2 expression). The hydrogel alone also exerted beneficial therapeutic effects, including reduced damage scores, increased Col2 expression, and reduced Mmp13 levels; however, these effects were notably smaller than those achieved with hiEV treatment while it was independent of the medium used for hiEV collection.

**Conclusions:** Together, our findings demonstrate that hiEVs from xeno-free conditions effectively prevent cartilage degradation and promote its repair. This paves the way for future clinical translation of hiEV-based therapies as a safe, scalable, and effective approach to treat osteoarthritis.

## Background

Osteoarthritis (OA) is a progressive and multifaceted joint disease which affects millions of individuals worldwide, especially in the aging population (1). OA pathophysiology is marked by degeneration of cartilage, remodelling of subchondral bone, and downstream processes frequently involve inflammatory responses of the synovium and other joint structures (2). Repairing damaged cartilage is particularly challenging due to its limited capacity for regeneration (3). Current OA treatments are limited to lifestyle changes, physical therapy, and medications aimed at symptom management whereas surgical options concern total joint arthroplasty (TJA) at end stage OA. Although TJA is currently considered the only disease modifying treatment, approximately 30% of patients remain dissatisfied with the outcomes (4). Consequently, there is an increasing interest in innovative therapeutic approaches to meet the growing needs of individuals affected by OA (5).

Human induced mesenchymal stromal cells (hiMSCs) derived from induced pluripotent stem cells (hiPSCs) offer a scalable and consistent source of stem cells for regenerative therapies. In addition to their scalability, hiMSCs exhibit greater proliferative potential and are less prone to donor-related variability as compared to isolated mesenchymal stromal cells (MSCs), making them particularly suitable for large-scale therapeutic applications (6). Even more, MSC-derived extracellular vesicles (EVs) have emerged as a promising cell-free therapy with significant regenerative potential for the treatment of OA (7, 8). EVs are small membrane-bound particles that are naturally released by cells and contain bioactive molecules such as proteins, lipids, RNA, and/or DNA. A proportion of EVs are involved in mediating intercellular communication, and if they originate from specific cells, such as MSCs, they can modulate pathophysiological processes to induce regeneration and tissue repair (6, 9). The culture conditions of MSCs, however, are suggested to critically influence the molecular and functional characteristics of EVs. In particular, serum-free and serum-containing media differentially affect EV yield, composition, biological activity, and overall therapeutic potential (10, 11). Nonetheless, EVs derived from MSCs have been shown to support cartilage regeneration by promoting chondrocyte proliferation, reducing inflammation, and enhancing production of extracellular matrix (ECM) (12, 13).

In a rat OA model, MSC-EVs improved cartilage repair in osteochondral defects However, producing large quantities of EVs for clinical applications is challenging because of the aging processes of primary MSCs and the intradonor heterogeneity of EV releasing MSCs, resulting in batch-to-batch variations in MSC-EV products (14). In this respect, hiMSC-derived EVs (hiEVs) can address these challenges and offer a more consistent, scalable source for cell-free OA therapy (15).

To explore the therapeutic potential of hiEVs for OA treatment, either in serum-containing or serum-free (PurStem) media (hiEV_Serum and hiEV_PS, respectively), we used a destabilizing medial meniscus (DMM) mouse model. Moreover, we explored potential advantages of a recently developed hydrogel as an effective delivery system of hiEVs for intra-articular (i.a.) injection while offering extended retention at the target site (16).

## Materials and methods

### Extracellular vesicle isolation and characterization

Generation and characterization of the hiMSCs used in this study has been described previously(17). HiMSCs were cultured under two different conditions. For serum-containing medium, hiMSCs were cultured in DMEM-GlutaMAX medium (GiBCo, Waltham, MA, United States) supplemented with 10% FBS (Sigma–Aldrich, St. Louis, MO, United States), 1 ng/ml fibroblast growth factor-2 (FGF-2, Peprotech, London, United Kingdom) and 100 U/ml penicillin-100 mg/ml streptomycin mixture (GiBCo). For the serum-free condition, the cells were continuously cultured in PurStem xeno-free medium (18) devoid of any serum contamination. The cells were maintained at 20% O2 and 5% CO_2_ at 37°C, and passaged upon reaching ∼80% confluence.

To isolate hiEVs from serum-containing and serum-free media, conditioned medium (CM) was collected from cell cultures after 72 h and immediately centrifuged at 300 × *g* for 10 min at 4°C to remove dead cells and debris. The supernatant was then centrifuged at 2000 × *g* for 20 min at 4°C to eliminate apoptotic bodies and stored at -80°C. For hiEV isolation by ultracentrifugation, CM from passages 7 to 11 was pooled and subjected to ultracentrifugation as previously described (17, 18).

Characterization of hiEVs was performed according to the MISEV criteria (19) including standards for surface markers CD9, CD63, and CD81 (**Supplementary Figure S1**). Data acquisition was performed by ImageStreamX Flow Cytometry (IFCM) for hiEV_serum, or by FACS on an Amnis ImageStreamX MkII instrument for hiEV_PS. Analysis and gating were carried out via IDEAS Software 6.2 according to the established workflow, as previously described, including the quantification of single EV events via custom fluorescence and spot count masks (20, 21).

Additionally, to follow the site of injection and check retention at the site of injection of the hiEVs in the mice, they were labeled with IR780 as previously described (22). Briefly, EVs were incubated with 100 μM IR780 at 4°C for 30 min and subsequently passed through a size exclusion purification column (Exo-spin®) to remove any unbound dye according to the manufacturer’s protocol. This step ensured that only labeled EVs were collected, increasing the specificity of the labeling.

### Measurement of hiEV release with degradation of the thermosensitive hydrogel

The release kinetics of hiEVs from hydrogels were determined via the direct release method (23, 24). Briefly, 1×10^9^ hiEVs were mixed with 1 mL of 25% (w/w) hydrogels in 15 mL tubes at 4°C. Afterwards, the tubes were incubated at 37°C for 5 min to allow the formation of a stable gel. Then, 4 mL of prewarmed PBS was gently added on top of the gel layer in the tubes, and the tubes were placed in a shaking water bath at 37°C and 30 rpm. At specified time points (1, 2, 4, 8, 12, 24, 48, 72, 96, 120, 144 and 168 h), 1 mL of the supernatant was collected from the tube, and the volume was replaced with fresh 37°C PBS. To determine the concentration of EVs in the supernatant, Nanoparticle tracking analysis (NTA) was performed via a Nanosight® NS300 (Malvern).

In addition, the degradation of the hydrogels was evaluated. To this end, 1 mL of hydrogel was loaded in 15 mL Falcon tubes and placed at 37°C for gel formation. The initial weight of the tube with the hydrogel was taken as 100% gel weight. Afterwards, 4 mL of prewarmed 37°C PBS was added on top of the hydrogel. At predetermined time points (corresponding to those in the release study), the entire volume of PBS was removed, and the change in weight of the conical tube with respect to the remaining hydrogel was measured to calculate the amount of hydrogel degradation.

### *In vivo* experiments

#### Mouse osteoarthritis model and experimental design

For the in vivo experiment, 40 male 12 week-old C57BL/6J mice were purchased from Charles River Laboratories (Charles River, Chatillon-sur-Chalaronne, France). The animal procedures were all conducted at the Leiden University Medical Center and were approved by the Animal Welfare Committee (IvD) under number AVD1160020171405-PE.18.101.006 and in line with ARRIVE guidelines 2.0. All mice were housed in groups in polypropylene cages (4 animals/cage) on a 12-hour light/dark cycle with unrestricted access to standard mouse food and water. The first group (N=4 mice) served as a sham (positive) control. The remaining 36 mice underwent surgery for destabilization of the medial meniscus (DMM) to establish a knee osteoarthritis model as described elsewhere (25, 26). Briefly, 30 min before surgery, animals pre-operative analgesia, Buprenorphine, were administered by Subcutaneous injection. The animal was then placed under isoflurane anaesthesia and the incision was started on the right knee when the animal no longer displayed reflex while the breathing was constant. A 1-cm longitudinal medial para-patellar incision was made to expose the knee joint, afterwards opened the knee joint gently through lateral dislocation of the patella and patellar ligament. The medial meniscotibial ligament was cut which anchors medial meniscus to the tibial plateau. After transection, the knee joint capsule was closed with a 6-0 absorbable suture and the skin closed with biological glue. Mice were immediately transferred to a heated post-operative recovery room. All animals received buprenorphine HCl (Vetergesic; Alstoe Animal Health, York, UK) sub-cutaneously post-surgery every 8h for the next 48h and were monitored daily to ensure good health. Successful destabilization of the medial meniscus was confirmed during surgery. Animals were then randomly allocated to experimental groups using dice. Intra-articular (i.a.) injections were performed by a colleague not involved in randomization, while the injection samples were prepared and coded by another colleague not involved in the animal experiments to ensure blinding. Twenty-one days after surgery, the DMM mice were treated with one time intra-articular injection of 5 µL PBS, 107 hiEVs from serum-containing media with/or hydrogel, or 10^7^ hiEVs from PurStem (PS) media with/or without hydrogel, as described in **Figure 1**. The mice were monitored daily to confirm their general health indicators according to their body weight and knee diameter and were sacrificed 35 days after the injection for analysis.

**Figure 1:**
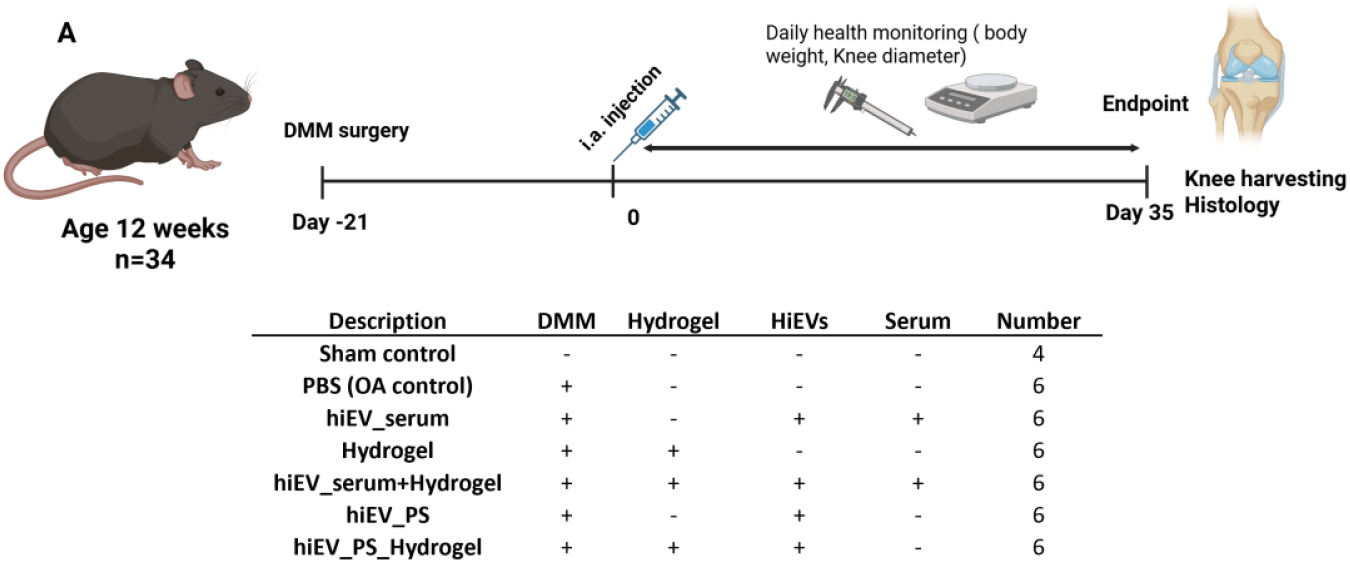
Schematic overview of the experimental timeline for *in vivo* treatment. 21 days prior to the treatment, OA was induced by performing DMM surgery followed by a single i.a. injection of hiEV_serum or hiEV_PS (± hydrogel), hydrogel alone, or PBS. Mice were monitored along the trajectory and sacrificed on day 35. DMM= Destabilization of the Medial Meniscus; i.a.= intra articular; OA= Osteoarthritis.

A Pearl Impulse Imaging System (LI-COR, Lincoln, NE, USA) was used to assess hiEV retention in the knee joint based on fluorescence. The mice were imaged at days 1, 7, 14, 21, 28, and 35 after i.a. injection under isoflurane anesthesia (4–5% induction, 1–2% for maintenance). Imaging was performed in the 800 nm NIR fluorescence channel (IR780 emission). The fluorescence intensity was quantified via Image Studio Software (LI-COR Biosciences), with a region of interest (ROI) around the knee joint. Background fluorescence was corrected using a nonfluorescent ROI.

### Histological evaluation

Following sacrifice of the mice, the limbs were collected for histological assessment. Knee joints were fixed for two days in 4% buffered paraformaldehyde, followed by 5 days of decalcification using a commercial mol-decalcifier (Milestone; pH 7.4) at 37 °C. The joints were then embedded in paraffin and sectioned at 5 μm thickness. Sections were stained with Safranin O/Fast Green and hematoxylin– eosin (H&E) and examined by light microscopy to evaluate cartilage damage in the medial compartment of the knee.

The commercial decalcification method was found to markedly reduce glycosaminoglycan (GAG) staining intensity compared with the standard 10% EDTA (pH 7.3) protocol, as observed with Safranin O, Toluidine Blue (TB), and Alcian Blue (AB) staining (**Supplementary Figure S2A**). To enable scoring despite this limitation, we developed a modified OARSI scoring system (damage OARSI score) based on direct visualization of structural cartilage damage in the cartilage zones using H&E and Safranin O staining as described in ‘Grading of cartilage’. This approach emphasizes parameters such as surface integrity, cartilage thickness, and chondrocyte organization (27) and was validated by scoring sections decalcified with 10% EDTA both with the modified damage OARSI scoring system and conventional OARSI scoring, and comparing the results with Spearmann correlation analysis (**Supplementary Figure S2B**).

### Grading of Cartilage

In DMM mouse models of OA, lesions are most severe in the medial compartment, which includes the medial femoral condyle (MFC) and medial tibial plateau (MTP). Joint degeneration was assessed using a cartilage damage score based on histological evaluation, following principles similar to the OARSI scoring system (25).

Representative grading criteria for the damage OARSI scoring system are shown in **Supplementary Figure S3**. Grading of the MFC and MTP in each section was performed by three independent researchers, blinded to experimental conditions and to each other’s scores. The results were averaged, and the mean damage OA score for each section was taken as the representative measure of knee joint damage, as described elsewhere (26).

### Immunohistochemical staining

Immunohistochemical (ICH) staining was performed on knee joint sections for collagen type II (Col2) and for matrix metalloproteinase (Mmp13). In brief, paraffin sections (5 µm) were blocked for endogenous peroxidase activity via incubation with 0.3% H_2_O_2_ for 10 min at room temperature. Antigen retrieval was performed with 25 µg/ml proteinase K (Prot K) prepared in 0.1 M Tris/HCL, pH 5.0, for 10 min at 37°C, followed by 30 min of treatment with hyaluronidase (5 mg/ml in Tris/HCL, pH 5.0). All the sections were blocked in 5% PBS-BSA for 30 min at room temperature. The following primary antibodies were used: anti-type II collagen (Col2) (Abcam, ab34712, Abcam, Cambridge, MA, dilution 1:200) and anti-matrix metalloproteinase (Mmp)-13 (sc-515284, Santa Cruz Biotechnology, Santa Cruz, Dallas, TX, USA, dilution 1:200). Primary antibody incubation was performed overnight at 4°C. Normal mouse IgG1 (2 µg/mL) served as an isotype control (sc-3877; Santa Cruz, Dallas, TX, USA). All the antibodies were diluted in 5% PBS-BSA. The next day, the slides were incubated with anti-mouse HRP (Envision, Dako, CA, USA) for 30 min at room temperature and subsequently incubated with a liquid DAB + 2-component system (Agilent, Santa Clara, CA, USA) for 5 min. The sections were counterstained with hematoxylin, dehydrated, cleared in xylene and covered with glass using Eukitt mounting medium (Sigma–Aldrich, Saint Louis, USA). Quantification of the IHC intensity was performed via ImageJ software as described elsewhere (28).

### Statistical analysis

Optimal sample size was determined based on statistical analysis of the results of similar DMM studies. For statistical analysis, GraphPad Prism 8.1.1 software (GraphPad Software, San Diego, CA, USA) was utilized. All data are expressed as the mean ± standard deviation (SD) of 3-5 independent repeated experiments, unless otherwise stated. Statistical significance was determined using Student’s t-test, unpaired, Mann-Whitney U test, and two-way analysis of variance (ANOVA). *P*-values were determined by applying a linear generalized estimating equation (GEE) via IBM SPSS statistics 25 to estimate the independent effects of covariables. In all analyses, a *P* -value≤0.05 is considered an indicator of statistical significance and is expressed as: * P≤0.05, ** P≤0.01, *** P≤0.001, **** P≤0.0001.

## Results

### Hydrogel enhances sustained EV release

hiEVs were collected and concentrated from conditioned media via ultracentrifugation, subsequently quantified by nanoparticle tracking analysis (NTA) and characterized before storage (**Supplementary Figure S1**). Before application in the *in vitro* release assay, hiEVs were incorporated into a thermosensitive hydrogel composed of 25% poloxamer 407 and 1% (w/v) self-assembling peptide (16) to assess whether the hydrogel could support the sustained release of hiEVs over time. Therefore, at predetermined time points, supernatants from IR780-labeled hiEVs mixed with hydrogel were collected and analyzed via NTA. The hydrogel release profile of hiEVs revealed an increase in fluorescence over time, indicating that progressive hiEV release closely mirrored the degradation of the hydrogel matrix over a 9-day period (**Figure 2A**).

**Figure 2.**
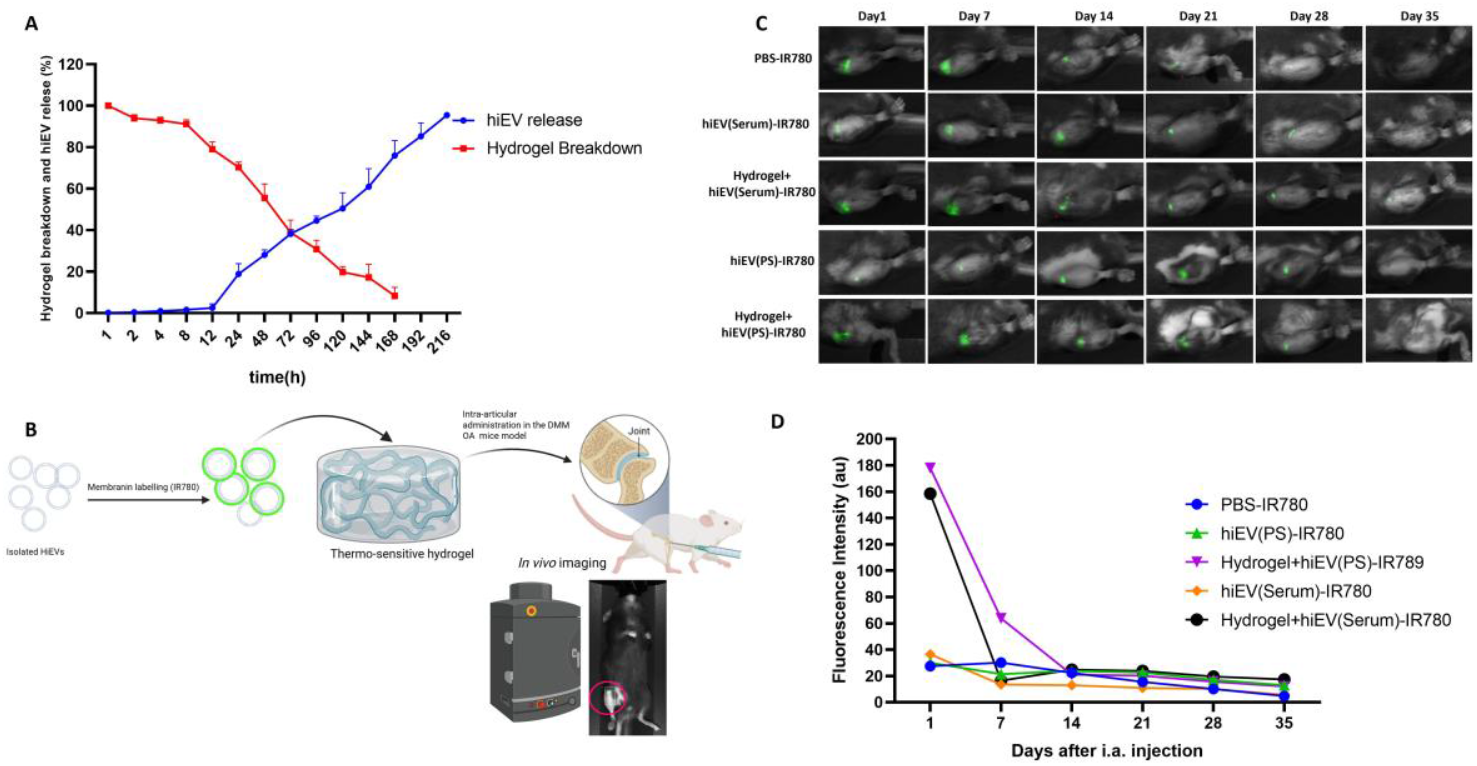
Sustained release and *in vivo* retention of hiEVs with or without thermosensitive hydrogel. (**A**) *In vitro* release profile of hiEVs and concurrent hydrogel degradation over 9 days at 37 °C. hiEVs were incorporated into a thermosensitive hydrogel composed of 25% poloxamer 407 and 1% (w/v) self-assembling peptide. EV-release was quantified by NTA, and hydrogel breakdown was assessed in parallel (n = 3 per time point; mean ± SD). (**B**) Schematic overview of *in vivo* experimental design. hiEVs were labeled with IR780, mixed with or without hydrogel (50% labeled, 50% unlabeled), and injected intra-articularly (10 × 106 hiEVs per knee) into DMM-operated mouse knees. Retention of labeled hiEVs was monitored weekly using near-infrared (NIR) fluorescence imaging. (**C**) Representative longitudinal fluorescence images of mouse knees at days 1, 7, 14, 21, 28, and 35 postinjection, showing signal retention across treatment groups. (**D**) Quantification of fluorescence intensity over time shows enhanced retention in the hydrogel+hiEV groups during the first 7 days, with comparable low signal levels across all groups by day 14 (n = 4–6 mice per group; mean ± SD). NTA= nanoparticle tracking analysis

Next, we assessed the release profiles of IR780-labeled hiEVs following intra-articular (i.a.) injection into the knee joints of OA mice (**Figure 2B**). PEARL Impulse imaging was used to monitor weekly the fluorescence intensity at the site of injection over the course of five weeks, and the results were compared across the different groups (**Figure 2C)**. hiEVs were administrered either alone or incorporated into the hydrogel. One day post-injection, the signal intensity markedly decreased in the groups without hydrogel, indicating rapid clearance of free hiEVs from the joint cavity. In contrast, hydrogel-embedded hiEVs retained strong fluorescence on day 1, suggesting improved short-term retention. By day 7, however, the fluorescence signals had decreased across all groups, and comparably low levels were detected from day 14 onward, as confirmed by signal quantification (**Figure 2D**).

### hiEVs slows cartilage degeneration in a DMM mouse model

Histological analysis was performed to evaluate joint damage across the experimental groups. To account for reduced proteoglycan staining observed with the commercial decalcification method, a modified damage OARSI score was applied, focusing on structural cartilage changes. Validation against standard OARSI scores from previous EDTA-based datasets demonstrated a strong positive correlation (*r* = 0.763, *P* < 0.01; **Supplementary Figure S2B**) substantiating the alternative approach.

Histological evaluation revealed that sham-operated mice exhibited minimal joint damage, whereas PBS-treated DMM mice presented pronounced cartilage degradation and damaged structure (**Figure 3 A-B**). In contrast, mice treated with hydrogel and hiEVs (+/-hydrogel) regardless of the culture medium, exhibited significantly reduced damage (**Figure 3A-B**).

**Figure 3.**
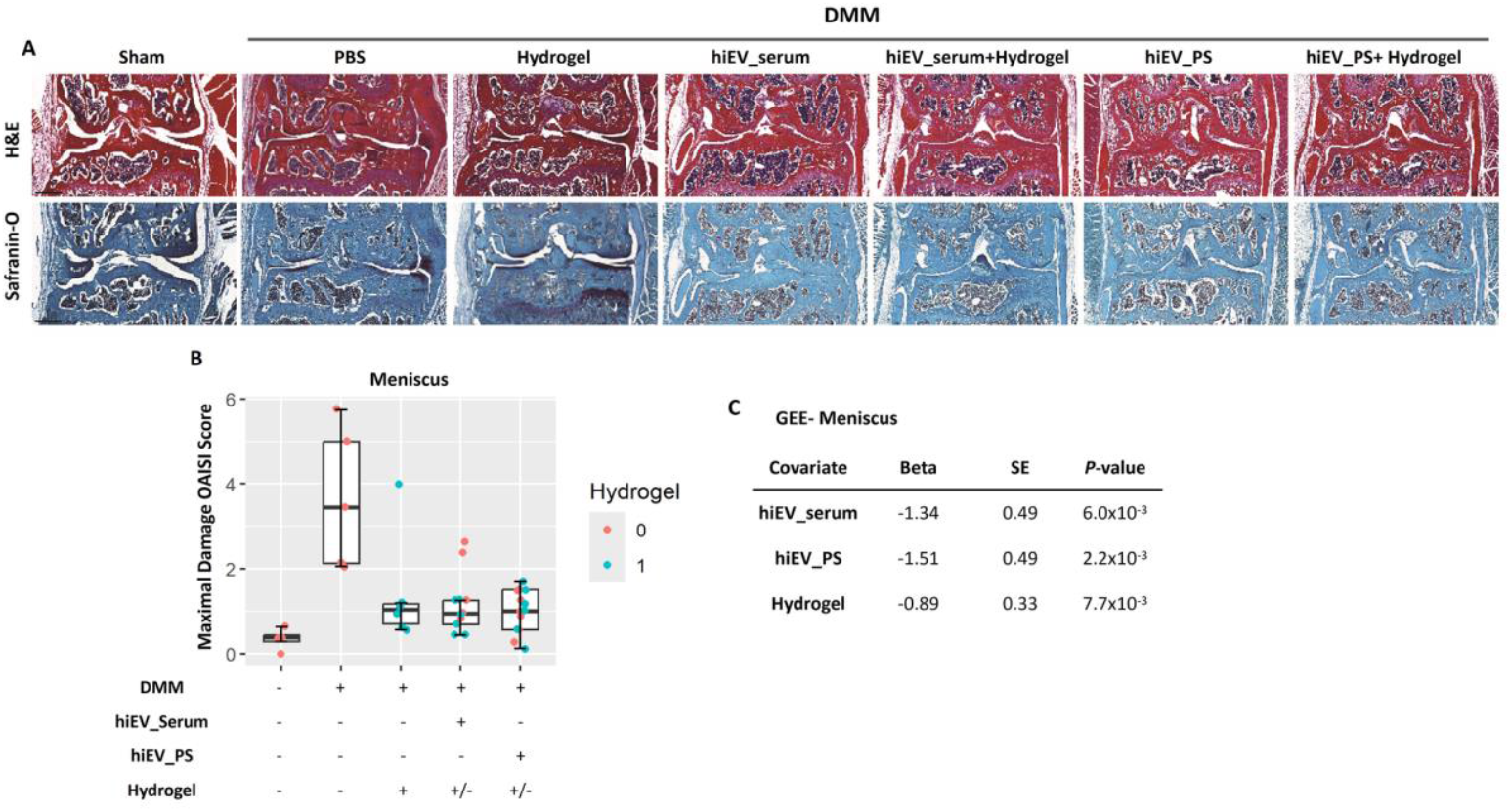
hiEVs reduce OA-associated joint damage *in vivo*. (**A**) Representative histological images of knee joints stained with H&E and Safranin-O/Fast Green from sham, PBS-injected DMM, and treated groups. (**B**) Quantification of cartilage damage scores in the meniscus area across treatment groups. Data are shown as the mean ± SD (*n* = 6). (**C**) Multivariate analyses of damage scoring using GEE to understand the independent therapeutic effect of. hiEVs derived from both serum and PS and hydrogel in the meniscus lesions. All data are shown as means ± standard deviations (n= 6). DMM= Destabilization of the Medial Meniscus, H&E= Hematoxylin and Eosin; i.a.= Intra-articular; GEE= Generalized Estimating Equation.

To independently evaluate the effects of, hiEV_serum, hiEV_PS, and hydrogel, a multivariate regression analysis using Generalized Estimating Equations (GEE) was performed. As shown in **Figure 3C**, all treatment groups exhibited significantly reduced damage scores relative to PBS-treated DMM mice. Specifically, treatment with hiEV_serum (Beta=–1.34, *P*=6.0×10^−3^) and hiEV_PS (Beta=–1.51, *P*=2.2×10^−3^) both resulted in significant reductions in cartilage damage. Hydrogel alone was also found to exert a significant chondroprotective effect, though this effect was smaller than that observed for hiEV-based treatments (Beta = -0.89 *P*=7.7×10^−3^).

Together, our results demonstrate that hiEVs exhibit robust protective effects against joint tissue degradation, with hydrogel delivery offering an additional, albeit smaller, therapeutic benefit.

### hiEV treatment reduces catabolic activity and enhances anabolic responses in a DMM mouse model

Following our observation that hiEVs with/or hydrogel exert therapeutic effects in the DMM model, we next evaluated their impact on catabolic and anabolic processes. We focused on matrix metalloproteinase 13 (Mmp13), a catabolic enzyme associated with cartilage degradation, and type II collagen (Col2), a key anabolic ECM component (**Figure 4A**).

**Figure 4.**
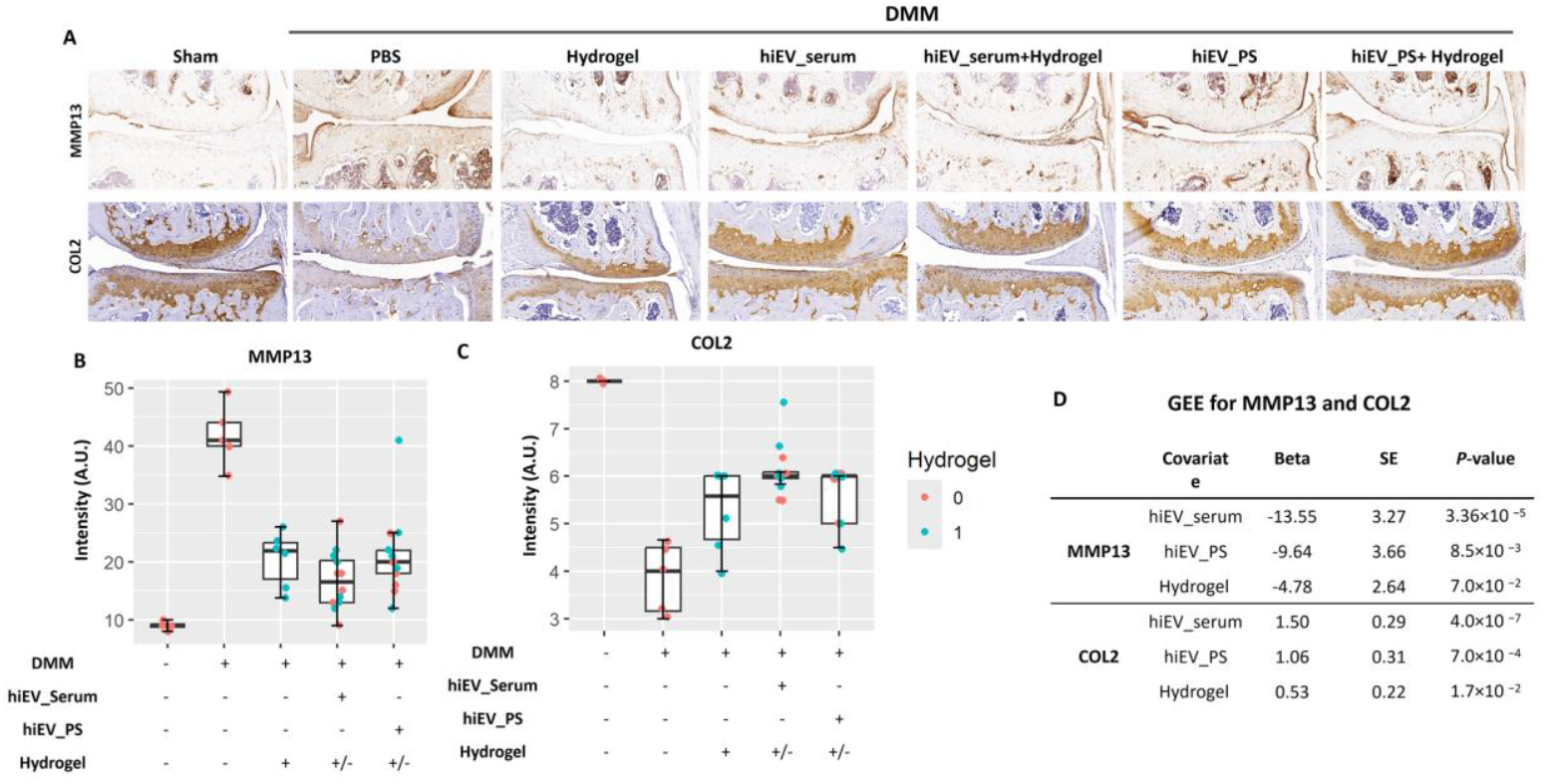
Immunohistochemistry analysis of Mmp13 and Col2 expression in the knee joint. **(A)** Representative image of mouse knees in the different treatment groups as indicated (Sham, DMM-PBS, Hydrogel, hiEV serum, hiEV serum + hydrogel, hiEV PS, and hiEV PS + hydrogel) stained for Mmp13 and Col2. **(B)** Quantification of IHC staining intensity for Mmp13 in the meniscus across the different treatment groups. **(C)** Quantification of IHC staining intensity for Col2 in the meniscus across the different treatment groups as indicated. **(D)** GEE analysis for Mmp13 and Col2 to understand the independent effect of hiEVs_serum and PS and hydrogel in the meniscus lesions. All data are shown as means ± standard deviations (n= 6 or 12). \*\**P*< 0.01 (0: no hydrogel; 1: with hydrogel). GEE: Generalized Estimating Equation.

Quantification of Mmp13 expression (**Figure 4B**) revealed the highest intensity values in PBS-treated DMM mice. Both hiEV treatment groups, with or without hydrogel, showed visibly lower expression levels, while hydrogel alone produced intermediate values between PBS and hiEV-treated joints. GEE analysis confirmed the independent effect of each treatment (**Figure 4D**). Mmp13 expression was significantly reduced in the hiEV_serum (Beta = –13.5, *P* = 3.4 × 10^−5^) and hiEV_PS (Beta = –9.64, *P* = 8.5 × 10^−3^) groups compared with PBS-treated DMM mice. Hydrogel alone also reduced Mmp13 expression, but the effect was not statistically significant (Beta = –4.77, *P* = 7.1 × 10^−2^).

**Figure 4C** shows quantification of Col2 expression across the different groups. PBS-treated DMM mice had the lowest values, whereas both hiEV treatment groups showed visibly higher Col2 levels. Hydrogel alone appeared to modestly increase Col2 expression relative to PBS. GEE analysis revealed significant increase in Col2 expression for hiEV_serum (Beta = 1.50, *P* = 4.0 × 10^−7^) and hiEV_PS (Beta = 1.06, *P* = 7.0 × 10^−4^) compared with PBS. Also hydrogel alone showed a slight improvement in Col2 intensity (Beta= 0.53, *P*= 7.0 × 10^−2^).

## Discussion

This study aimed to evaluate the therapeutic potential of human-induced mesenchymal stromal cell-derived EVs in an *in vivo* DMM-induced OA mouse model. Our findings confirmed that hiEV treatment showed protective effects against cartilage degeneration by promoting anabolic activity and suppressing catabolic pathways within the joint. Notably, the therapeutic efficacy of hiEVs was consistent regardless of the culture medium used for their production, whether serum-containing or xeno-free, highlighting the robustness and translational relevance of the use of hiMSC-derived EVs. In addition, the hydrogel delivery system proved capable of sustaining local hiEV availability within the joint cavity, thereby potentially enhancing their therapeutic effect.

Histological damage scoring revealed that hiEV treatment significantly reduced cartilage damage, particularly in the meniscal area of DMM mice. Because the commercial decalcification method used in this study markedly reduced proteoglycan staining, we modified the OARSI scoring system to focus on structural cartilage parameters visible in H&E and Safranin O staining. However, it may underestimate early degenerative changes where proteoglycan loss precedes structural breakdown. Therefore, our results primarily reflect structural preservation, while subtle biochemical changes might be under-represented. Additionally, IHC analysis showed upregulation of the anabolic marker Col2 and downregulation of the catabolic marker Mmp13, providing further evidence that hiEVs support extracellular matrix (ECM) preservation. This effect was comparable for hiEVs produced in serum-containing and xeno-free media.

The role of the culture environment in shaping EV properties is an ongoing subject of investigation. It has been suggested that serum components may adsorb onto EVs or become incorporated into their structure, potentially altering their composition and downstream bioactivity (13). In this context, the use of xeno-free media becomes particularly important when transitioning toward clinical applications, as it eliminates the risk of xenogeneic contamination and reduces the potential for immunogenic reactions (18–20). However, xeno-free production clearly offers advantages for regulatory approval and patient safety, reinforcing its relevance for future clinical application(29). These findings are also aligned with recent studies using iPSC-derived EVs produced under serum-free conditions, which similarly demonstrated cartilage-protective effects (30).

Notably, weekly monitoring of IR780-labeled hiEVs demonstrated their retention at the injection site, with no evidence of off-target distribution supporting both the safety and efficiency of the delivery system. This confirms the added value of the hydrogel delivery system in the absence of damaging effects, being localized delivery, facilitated sustained release, and simplified intra-articular application. Furthermore, the modest independent therapeutic effects of the hydrogel reflected by reduced damage scores and increased Col2 expression, suggest that it may provide a supportive microenvironment for cartilage preservation. However, the magnitude of these effects was notably smaller than that observed with hiEV treatment, underscoring that the primary benefit of the hydrogel lies in its role as a delivery vehicle. Similar strategies have been reported using thermosensitive polymers, such as poly (D,L-lactide)-b-poly(ethylene glycol)-b-poly(D,L-lactide) (PLEL) to successfully deliver MSC-EVs intra-articularly, demonstrating effective cartilage protection in rat OA models (31). Other injectable hydrogel systems, such as gelatin methacrylate (GelMA) combined with gelatin or nanoclay, and chitosan-hyaluronic acid blends, have also been utilized to enhance EV delivery and retention within joint tissues (32, 33). Our thermosensitive hydrogel offers comparable advantages of *in situ* gelation and minimally invasive application, while ensuring controlled and prolonged intra-articular availability of hiEVs.

Compared to their parental hiMSCs, hiEVs offer notable advantages including improved storage stability, lower immunogenicity, and reduced tumorigenic risk. The hiEVs can be manufactured and standardized more readily, supporting their use as a scalable and off-the-shelf therapy (34, 35). Although several studies have demonstrated that EVs have many of the same therapeutic effects as MSCs, some findings suggest that MSCs may exert broader or more sustained effects in certain contexts due to their ability to continuously secrete a wide range of bioactive factors,including cytokines, chemokines, and growth factors, not fully represented in EVs alone (36). To enhance the efficacy of hiEVs, future strategies should focus on optimizing EV loading and cargo composition, improving targeting and uptake efficiency, and potentially combining EVs with complementary biologics or scaffolds to recapitulate the multifaceted benefits of live cell therapies.

In conclusion, we provide compelling evidence that hiEVs produced under xeno-free conditions retain full therapeutic efficacy and can modulate both catabolic and anabolic processes in osteoarthritic cartilage. Combined with the practical benefits of hydrogel-based delivery, these findings pave the way for future clinical translation of hiEV-based therapies as a safe, scalable, and effective approach to treat osteoarthritis.

## Abbreviation

OA: Osteoarthritis
MSCs: Mesenchymal stromall cells
TJA: Total joint arthroplasty
EVs: Extracellular vesicles
hiMSCs: Human-induced Mesenchymal stromall cells
hiPSCs: human induced pluripotent stem cells
hiEVs: Extracellular Vesicle derived from Human-induced Mesenchymal stromall cells
ECM: Extracellular Matrix
PS: PureStem
DMM: Destabilization of the Medial Meniscus
IHC: Immunohistochemistry
Col2: Collagen Type II
Mmp13: Matrix Metalloproteinase 13
P407: Poloxamer 407
H&E: Hematoxylin and Eosin
NTA: Nanoparticle Tracking Analysis
I.A.: Intra-articular
GEE: Generalized Estimating Equations

## Ethics approval and consent to participate

The project “Dose finding & potential adverse effects observations in the DMM model” was conducted at the Leiden University Medical Center and was approved by the Animal Welfare Committee (IvD) under number AVD1160020171405-PE.18.101.006 on Apr 21, 2023.

The authors declare that they have not use AI-generated work in this manuscript.

## Consent for publication

Not applicable.

## Authors’ contributions

Sana S. Sayedipour: Collection and/or assembly of data (*in vivo* experiments), data analysis and interpretation, manuscript writing, final approval of manuscript.

Jelle Nikkels: Collection and/or assembly of data (*in vivo* experiments), final approval of manuscript. Tobias Tertel: Collection and/or assembly of data, final approval of manuscript.

Eka Suchiman: Collection and/or assembly of data, final approval of manuscript. Matilda Balbi: Collection and/or assembly of data, final approval of manuscript. Georgina Shaw: Collection and/or assembly of data, final approval of manuscript. Luis Cruz: Collection and/or assembly of data, final approval of manuscript.

Louise van der Weerd: Collection and/or assembly of data, final approval of manuscript.

Chiara Gentili: Collection and/or assembly of data, final approval of manuscript. Mary Murphy: Conception and design, final approval of manuscript.

Bernd Giebel: Conception and design, final approval of manuscript.

Ingrid Meulenbelt: Conception and design, data analysis and interpretation, manuscript writing, final approval of manuscript.

Yolande FM Ramos: Conception and design, collection and/or assembly of data, data analysis and interpretation, manuscript writing, final approval of manuscript.

## Acknowledgements

We sincerely thank the members of the OA research group of LUMC for their valuable feedback, scientific discussions, and continued support throughout this study.

## SUPPLEMENTARY MATERIAL

### Supplementary Figures

**Supplementary Figure S1.**
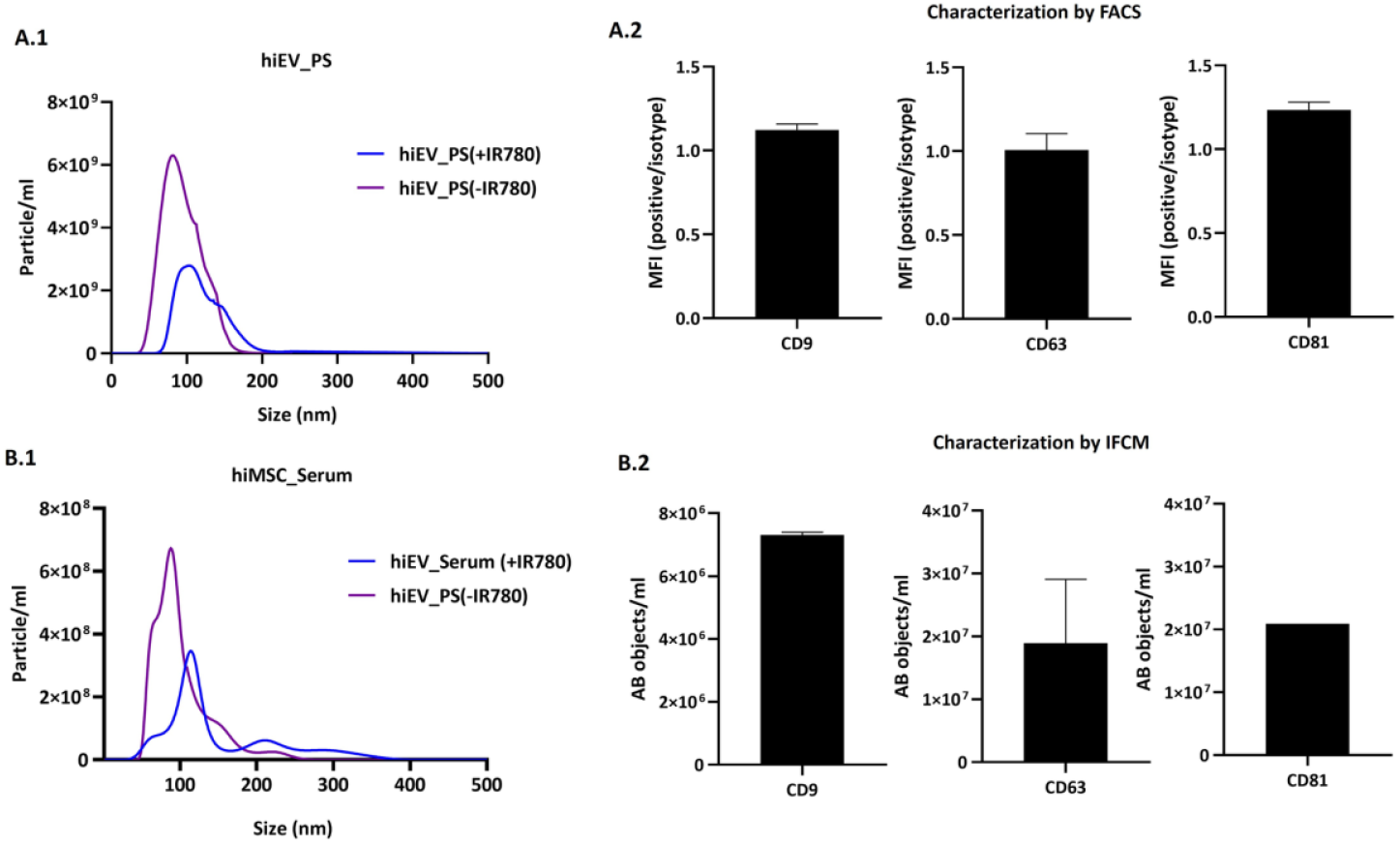
EVs characterization. A. hiEV PureStem (PS): **A1** NTA size distribution analysis of hiEVs isolated from SD1 P12 (n=3). **A2** Quantification of non conventional flow cytometry analysis: CD9-, CD63- and CD81-positive events in CFDA-SE gate. Data are shown as ratio between mean fluorescence intensity (MFI) of EVs stained with the specific antibody and the correspondent isotype control (relative MFI) (n=3). **B. hiEV_Serum**-containing: **B1** NTA size distribution analysis of hiEVs isolated from Serum containing medium (n=3). **B2)** Imaging flow cytometry (IFCM) results showing the concentration of objects/mL for CD9, CD63, and CD81.

**Supplementary Figure S2.**
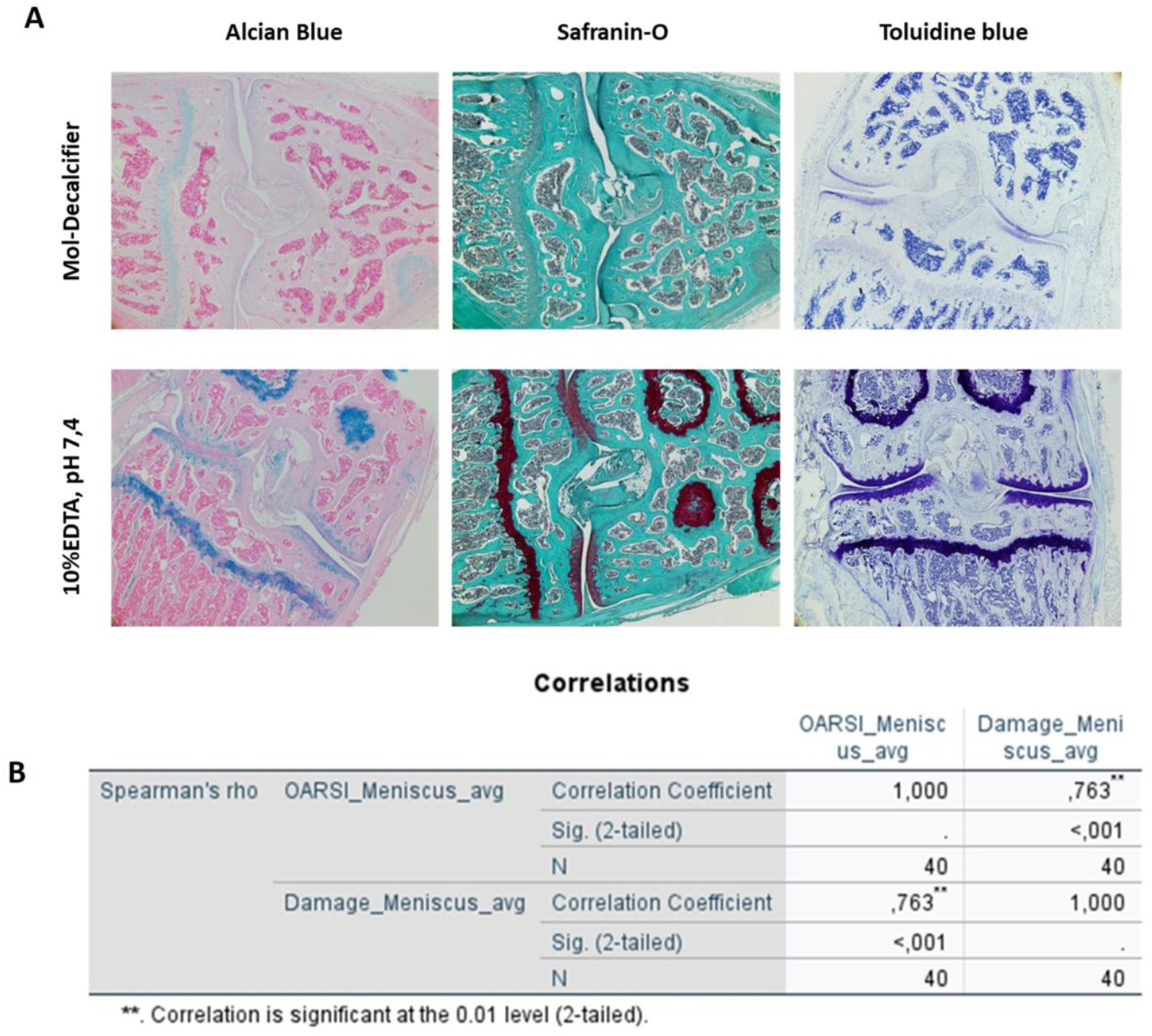
(A) Comparison of cartilage staining results using Mol-decalcifier and 10% EDTA (pH 7.4) for decalcification in DMM joints of OA mice samples. Staining with Alcian Blue, Safranin-O, and Toluidine Blue shows that the Mol-decalcifier method impacts glycosaminoglycan (GAG) visualization compared to the traditional EDTA method, with weaker GAG staining observed in Mol-decalcifier-treated samples. **(B) Correlation analysis between the newly developed damage scoring system and traditional OARSI scoring in meniscal cartilage**. Spearman’s correlation (r = 0.763, *P* < 0.01) demonstrates a significant positive relationship, indicating that the damage scoring system is a reliable and comparable alternative to the OARSI scoring system.

**Supplementary Figure S3.**
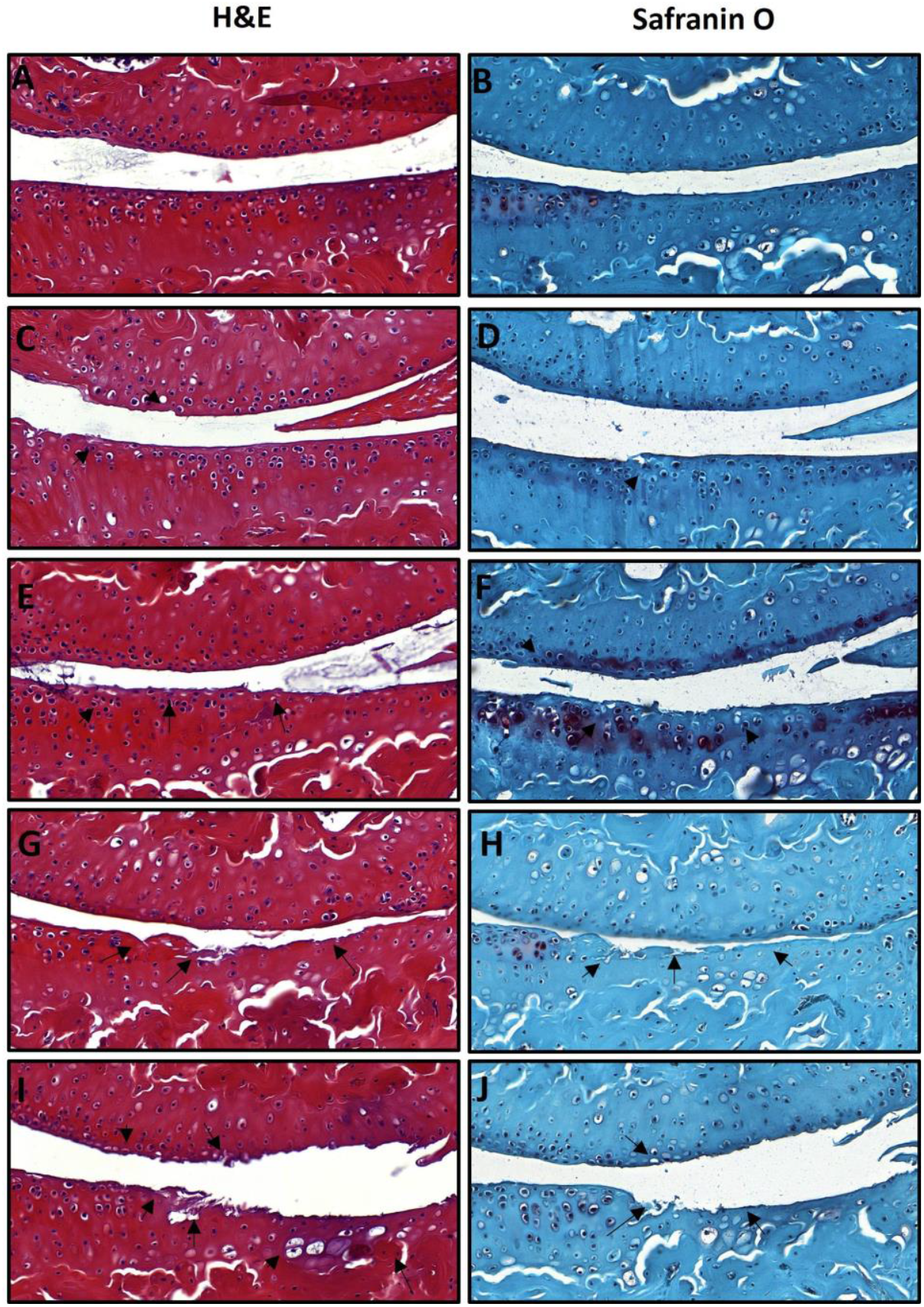
Histologic comparison of hematoxylin & eosin (H&E) staining and Safranin O staining of the articular cartilage with different grades of osteoarthritis. (**A)** and (**B**) The medial tibial plateau of a sham mouse with intact articular cartilage and grades based on the Damage OARSI schemes = 0. (**C**) and (**D**) The medial tibial plateau of a DMM mouse with Damage OARSI grade = 1. (**E**) and (**F**) The medial tibial plateau of a mouse after DMM with an Damage OARSI grade = 2-3. (**G**) and (**H**) The medial tibial plateau of a DMM mouse with Damage OARSI grade =4-5. (**I**) and (**J**) The medial tibial plateau of a DMM mouse with Damage OARSI grade =5-6

